# Neurological Tissue Dissection Techniques in Mouse Models for Reproducible Scientific Results: Brain, Spinal Cord, CSF, and Sciatic Nerve

**DOI:** 10.1101/2025.07.01.662633

**Authors:** Lilia Crew, Alyssa Seerley, Serena McElroy, Andrea Grindeland Panter

## Abstract

Biomedical research studies, specifically regarding human neurodegenerative diseases, are bound by ethical challenges, and have limited diagnostic and treatment options. Transgenic mouse models offer an incredible research advantage to conduct feasible and practical research with the ability to precisely define the progression of neurodegenerative disease over a wellcontrolled dosage and timeline. The use of transgenic mouse models has been extensive and is critical to advancing research in many ways, including understanding brain morphology and general tissue changes caused by neurological diseases. Often, these studies require specific brain regions or other neurological tissues which may be difficult to obtain. Unfortunately, specific extraction and dissection protocols are few and far between, leading to inconsistent results and a lack of reproducibility. A well-defined protocol, such as this, is instrumental in overcoming these obstacles and acquiring better experimental results. Five mouse-specific protocols are described: brain extraction, brain microdissection, spinal cord extrusion, cerebral spinal fluid (CSF) collection, and sciatic nerve dissection. Each protocol was completed under biosafety level 2 (BSL-2) guidelines, similar to the sterility precautions required in human surgery. Each protocol also includes a collective materials list that defines proper instruments and usage. The protocol was refined based on feedback from numerous research studies in transcriptomics and pharmaceutical development. These applications require minimizing tissue damage, dissection accuracy, and the ability to reproduce the results—skills that are also directly transferable to clinical settings. The proper implementation of these protocols will allow for more accurate and precise results with reduced variability. This study provides well-defined, succinct, accessible protocols that are more ethical and improve the overall quality of the conducted research. By addressing this need, it supports greater advancements in many cross disciplinary areas.

## 1. Introduction

Biomedical investigations into human neurodegenerative diseases face complex ethical challenges and are constrained by limitations of diagnostic and therapeutic modalities. Animal models have been used to study these diseases since the late 1950s, leading to revolutionary breakthroughs in the field (Young, 2009). Mouse models are particularly advantageous in understanding neurodegenerative diseases due to the remarkable similarity of the laboratory mouse, *Mus musculus*, to the human genome. Approximately 90% of these genomes can be divided into regions of conserved synteny, demonstrating the value and applicability of mouse models in the development of human medicine (Chinwalla *et al.*, 2002). Humans and mice share numerous developmental milestones at both structural and cellular levels, as well as exhibiting similar age-related behavioral traits (Tello *et al.*, 2022). Additionally, studies conducting an analysis of transcriptome networks in human and mouse brains have demonstrated significant conservation of gene expression and connectivity, as well as the preservation of network modules (Tello *et al.*, 2022).

Tissues in transgenic mouse models recapitulating neurodegenerative diseases are exceedingly valuable because experiments can be planned to have sample numbers which are statistically powered under controlled laboratory conditions, resulting in higher quality data and thorough comprehension (Ament *et al.*, 2017; Krakauer *et al.*, 2017; Voelkl *et al.*, 2018; Meyerhoff *et al.*, 2021; Cook, Hensley-McBain and Grindeland, 2023). What’s more, these tissues are practical to obtain through dissection and have been useful in studies of neurogenerative diseases such as transcriptomics, pharmaceutical development, and transgenic model development (Ament *et al.*, 2017; Minikel *et al.*, 2020; Ratz-Mitchem *et al.*, 2023; Bragg *et al.*, 2024). For example, the brain can be extracted and subsequently dissected into specialized regions to be used in disease studies that focus on specific regions of the brain. Additionally, biomarkers found in the brain and cerebrospinal fluid (CSF) have proven to be necessary in studies on various neurodegenerative disease models aiming to find diagnostic tests or pharmaceutical treatments (Kuttner-Hirshler *et al.*, 2017; Bondulich *et al.*, 2024).

Despite the value of these neurological tissues, dissecting them may present a multitude of challenges for researchers which may include maintenance of tissue integrity, efficient dissection time frames, obtaining specific tissues without contamination from other tissues, equipment limitations, and highly specialized training requirements. Brain tissue is extremely delicate and malleable; improper handling and omission of care can easily distort target structures. This can impact downstream analysis of histology and molecular assays, creating unreliable results that are hard to replicate (Meyerhoff *et al.*, 2021). Researchers also must be mindful of ischemia and tissue degradation in neurological tissue studies. A prompt extraction and proper fixation of tissues can help to eliminate the compromising effects of enzymes; however, it can require precise and technical skills. Without refined anatomical understanding, microdissection of specific tissues can be compromised, leading to inconsistencies in data and a lack of reproducibility across studies (Sultan, 2013; Meyerhoff *et al.*, 2021).

Various methods of mouse brain extraction and dissection have been previously reported (Liu and Duff, 2008; Sultan, 2013; Bala *et al.*, 2014; Richner *et al.*, 2017; Lim *et al.*, 2018; Meyerhoff *et al.*, 2021; Shimizu *et al.*, 2022; Kaur *et al.*, 2023; Aboghazleh *et al.*, 2024; Sell, Shi and Bhat, 2024), however our expanded methods have been refined with the dissection accuracy validated in studies across several disciplines (Ament *et al.*, 2017; Minikel *et al.*, 2020; RatzMitchem *et al.*, 2023; Vallabh *et al.*, 2023; Bragg *et al.*, 2024), are included together in a concise report, and do not require highly specialized equipment. Our well-defined protocols to maintain tissue integrity and experimental reproducibility are critical to correctly interpret experimental results, which in many cases, have taken years to acquire by aging animal neurodegeneration studies (Sultan, 2013; Aboghazleh *et al.*, 2024). These protocols will also enable the use of fewer research animals by increasing the sample quality and minimizing tissue loss (Voelkl *et al.*, 2018; Aboghazleh *et al.*, 2024).

## 2. Materials and Methods

### 2.1 Animals

The mice included in this study were housed in the Weissman Hood Institute at Touro University; McLaughlin Research Institute’s Animal Resource Center, which is an all-mouse facility and is accredited by the American Association for Accreditation of Laboratory Animal Care (AALAC). All animal studies were conducted in accordance with the US National Research Council’s Guide for the Care and Use of Laboratory Animals, the US Public Health Service’s Policy on Humane Care and Use of Laboratory Animals, and the Guide for the Care and Use of Laboratory Animals. The animal procedures and studies were reviewed and approved by the McLaughlin Research Institute’s Institutional Animal Care and Use Committee (IACUC).

### 2.2 Surgery Preparation

Before beginning any of these procedures, implement the appropriate biosafety level methods for proper safety and decontamination purposes, and familiarize yourself with an overview of procedural order (**Figure 1**). All instruments must be gathered as shown in **Figure 2** and sterilized in advance. CSF collection is the only procedure in this report performed under sedation as opposed to post-euthanasia. Administer the IACUC approved anesthesia if CSF collection will be performed. Ensure that the appropriate level of anesthesia is reached by lack of deep pain reflex and minimal respiratory response to a toe pinch with a hemostat. Following CSF collection, euthanize the mouse in accordance with AVMA and IACUC guidelines.

**Figure 1.**
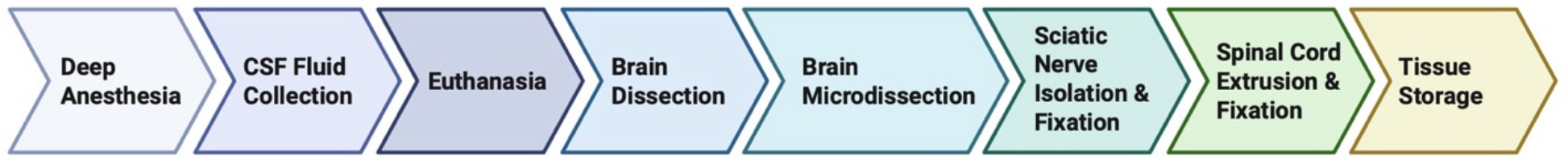
Experimental Workflow for CNS & PNS Dissection. Graphical overview of the sequential workflow highlighting the key stages of the dissection protocol. While shown in sequence, the protocol is modular, allowing researchers to adapt or omit steps based on specific experimental needs. Created in BioRender. Seerley, A. (2025) https://BioRender.com/rfi9rxe

**Figure 2.**
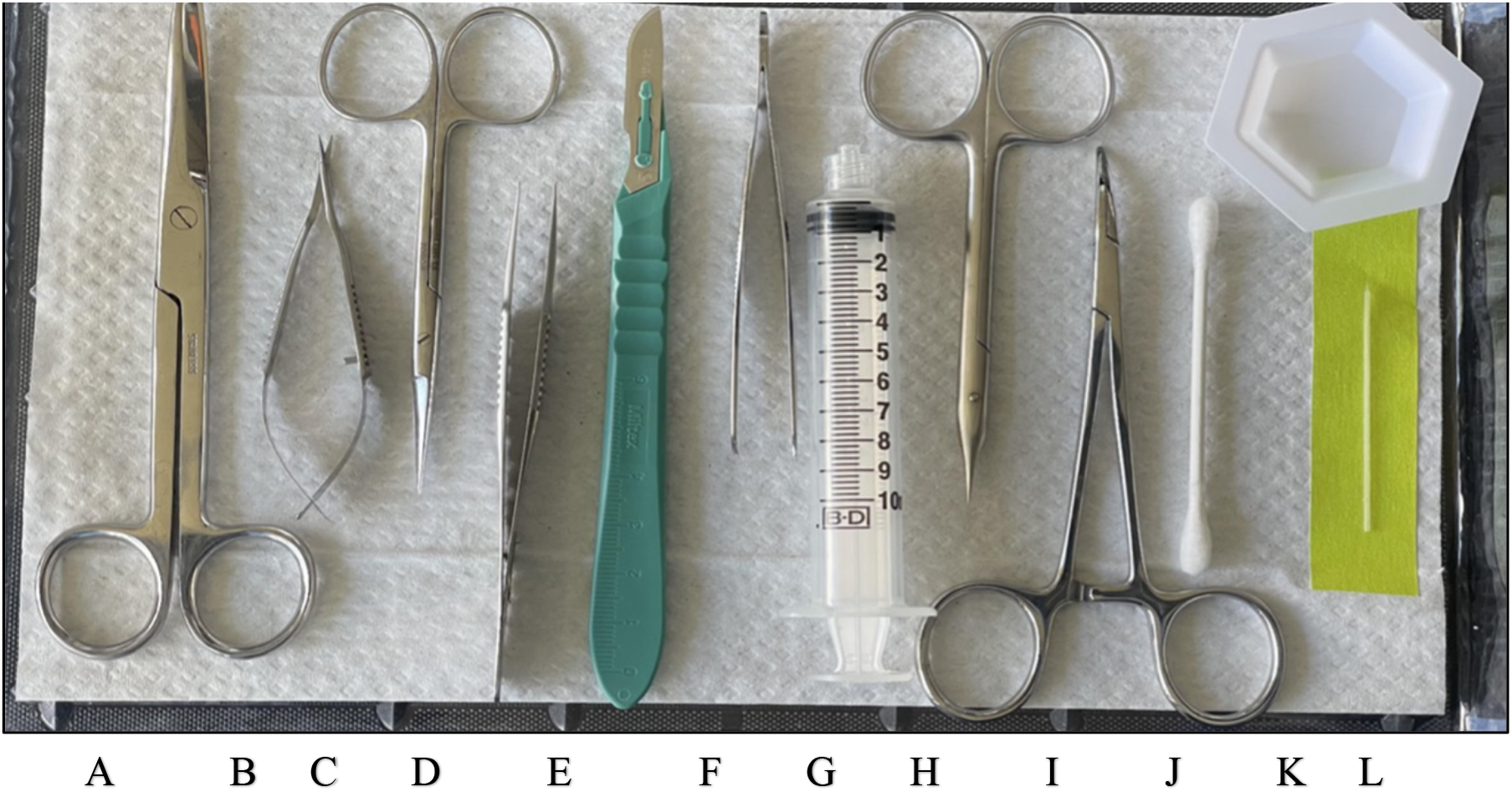
Dissection Instruments and supplies useful in microdissections. **[A]** Large Scissors; **[B]** Ophthalmic Microdissection Scissors; [**C]** Small Scissors; **[D]** Fine Tip Forceps; **[E]** Scalpel (#10 blade); **[F]** Curved Forceps; **[G]** Luer Lock Syringe (10mL); **[H]** Small Fine Tip Microdissecting Scissors; **[I]** Curved Hemostat, **[J]** Cotton Swab, **[K]** Weigh Boat; **[L]** Capillary Tube (on yellow tape). Not pictured: 70% EtOH, Petri Dish, PBS (1X concentration), Formalin (10%), Dissection Pins

### 2.3 Capillary Tube for CSF Collection

Using a micropipette puller, make a needle from a 10 cm length capillary tube made of polished borosilicate glass with filament, size O.D.:1.0mm I.D.: 0.78mm. The end of the needle will need to be cut to a diameter of 0.2mm, as the larger diameter will allow the CSF to flow into the tube by capillary action, see **Figure 3**.

**Figure 3.**
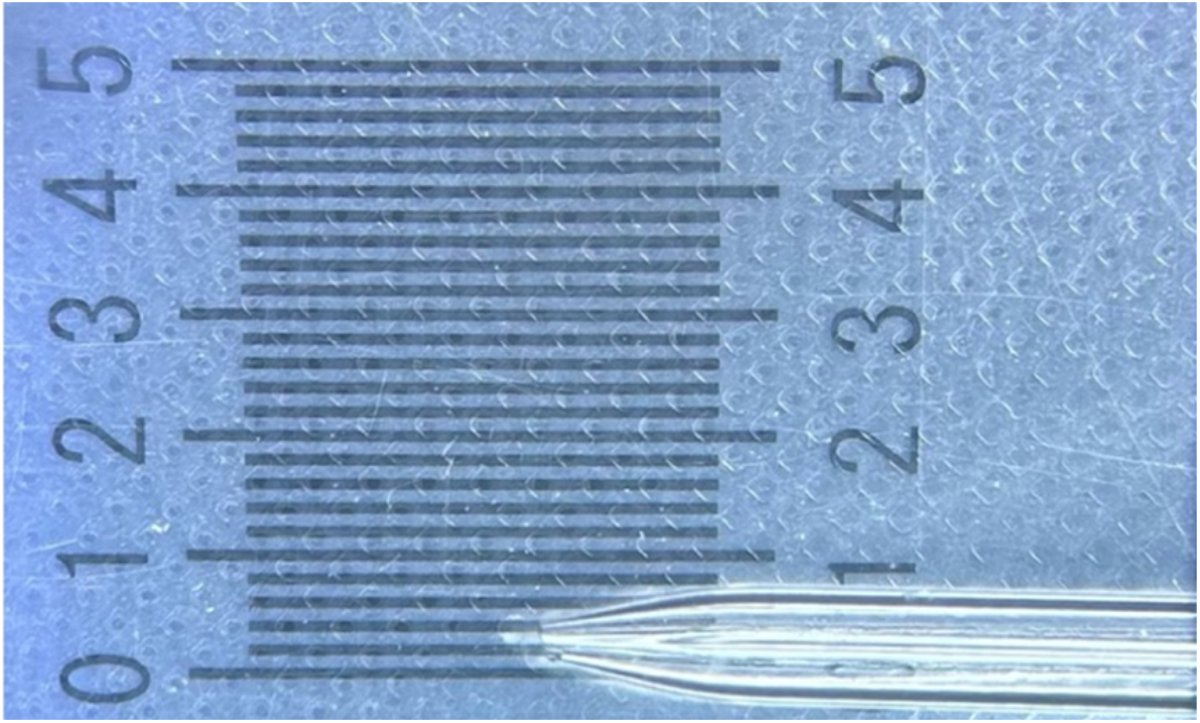
Capillary tube recommendations. Borosilicate glass with filament [O.D.: 1.0mm, I.D.: 0.78mm, 10cm length] capillary tube pulled into a needle and cut leaving a 0.2mm diameter opening for puncture of the arachidonic membrane.

### 2.4 Terminal Cerebral Spinal Fluid (CSF) Collection

1. Apply 70% ethanol (EtOH) to the surgical area to wet down the hair of the mouse.
2. Using a #10 blade, make a midline skin incision over the occiput to the second cervical vertebrae (C2).
3. Incise between the cervical muscles on the midline over the occiput to C2, using sharp dissection. The cervical muscles rhomboideus cervicis, the cervical part of the trapezius, splenius capitis, semispinalis capitis, and erector spinae muscles will all need to be separated at midline to expose the cisterna magna, however, the deeper muscles which are difficult to visualize with the eye may be separated under the dissecting microscope in step 6 using blunt dissection techniques.
4. Place the ventral surface of the thorax of the mouse on the weight boat under the microscope.
5. Stabilize the mouse ensuring that the skull is hyperflexed for maximum access to the cisterna magna, see **Figure 4**.
6. Bluntly dissect and separate at midline the interior muscles remaining from step 3 down to the cisterna magna to visualize a transparent membrane. This is the arachnoid membrane covering the cisterna magna, which is a large pocket of CSF within the subarachnoid space. If hemorrhage from the musculature occurs, blot it with a cotton swab prior to puncturing the arachnoid membrane as it is critical to avoid blood contamination in the CSF.
7. Once the arachnoid membrane is visualized, use small thumb forceps to move the muscles laterally if needed for complete visualization.
8. With the capillary tube, puncture the arachnoid membrane at a 45° angle, using a gentle spinning motion.
9. Allow CSF to flow into the tube by capillary action. The approximate amount of CSF collected is between 4-10µL, see **Figure 5**.
10. Place the blunt end of the capillary tube into the rubber bulb capillary dispenser instrument and expel sample into desired sample tube, see **Figure 6**.

**Figure 4.**
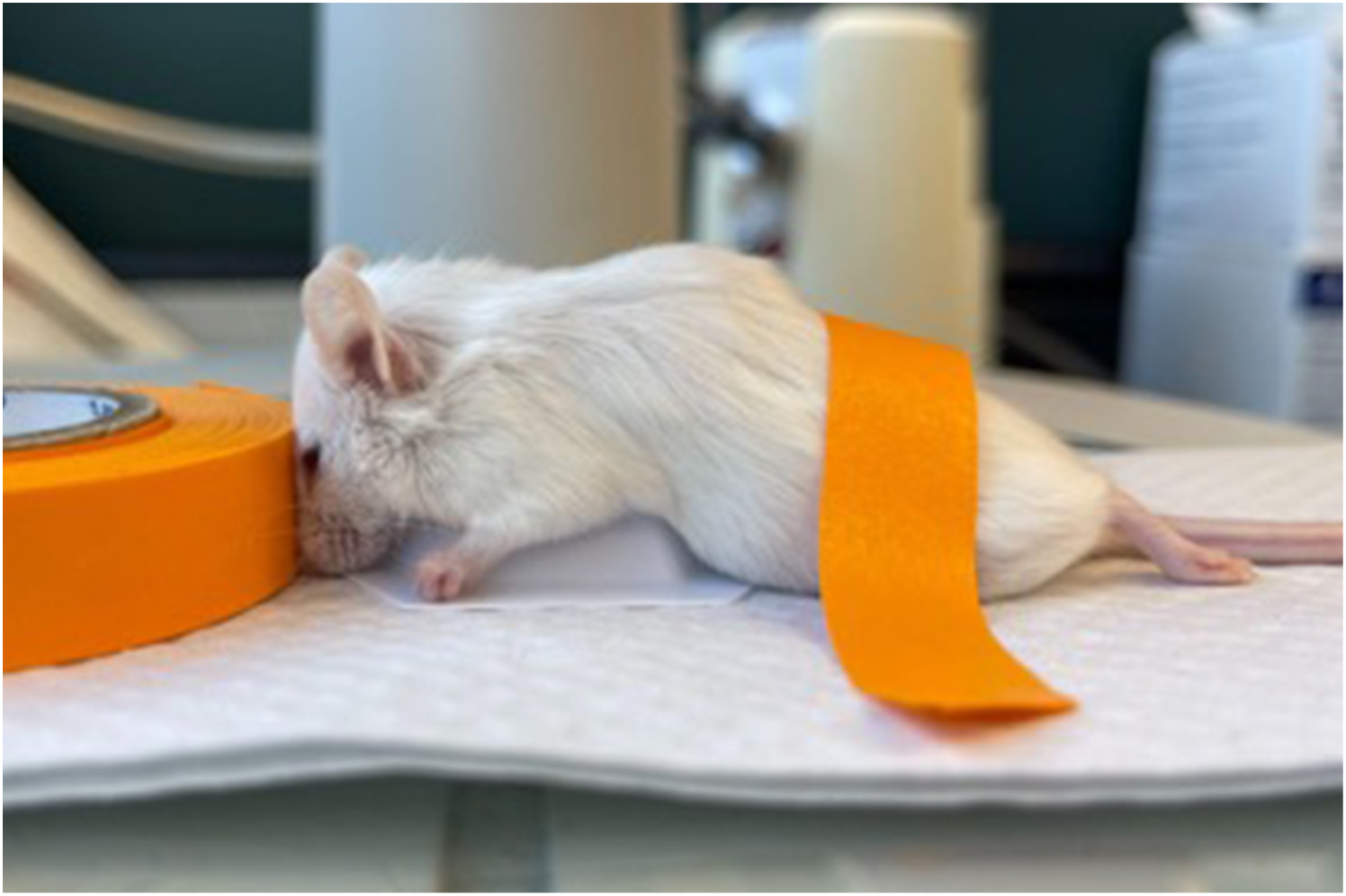
Mouse Positioning to obtain CSF. Anesthetized mouse positioned for cerebrospinal fluid (CSF) collection, demonstrating hyperflexion in the cranial-occipital region. The mouse is secured with orange tape to maintain the necessary posture for effective CSF extraction and hyperflexion is induced via a weigh boat and tape roll or any object that will hold the correct stabilization and placement of the head.

**Figure 5.**
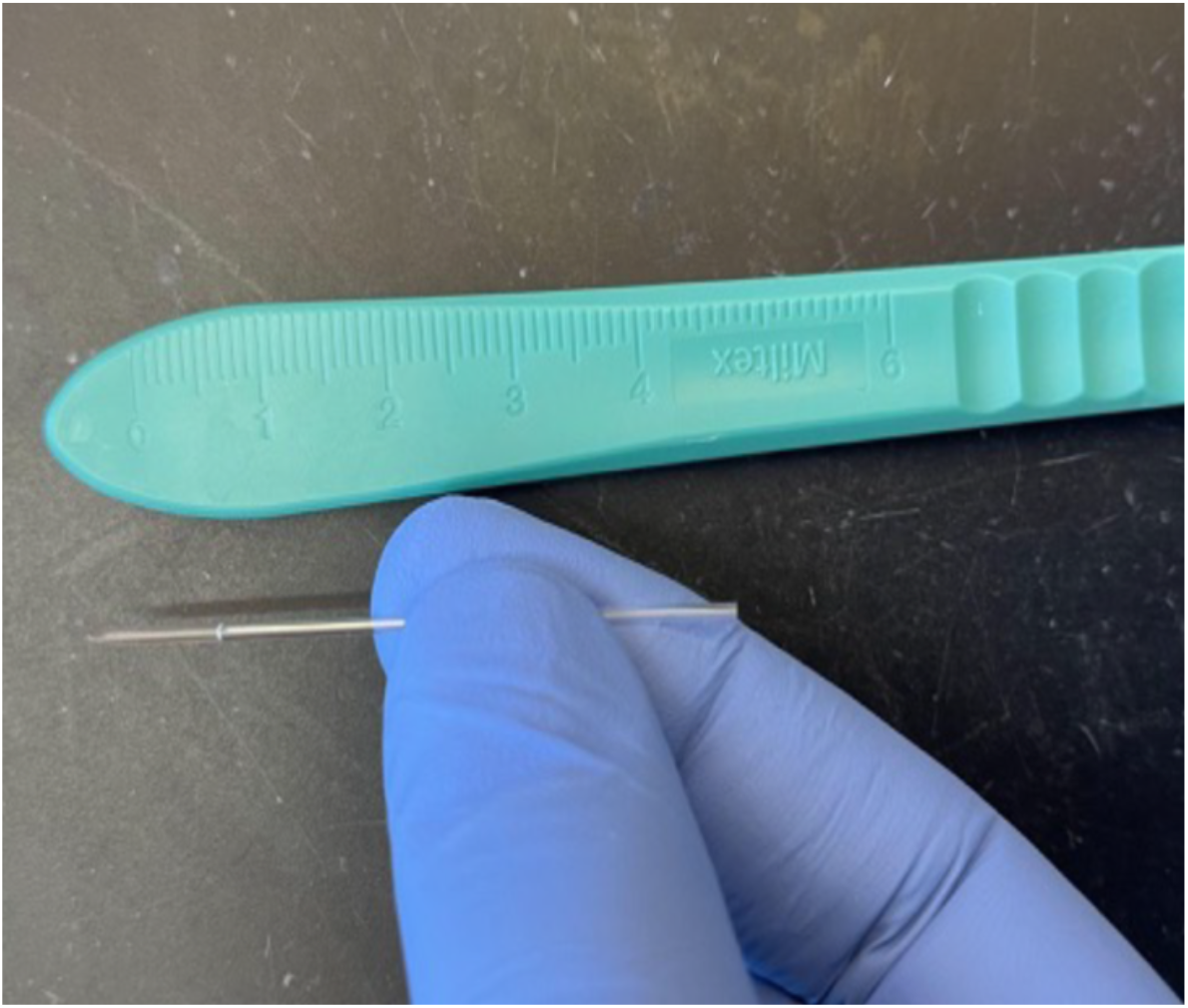
Capillary tube containing CSF.

**Figure 6.**
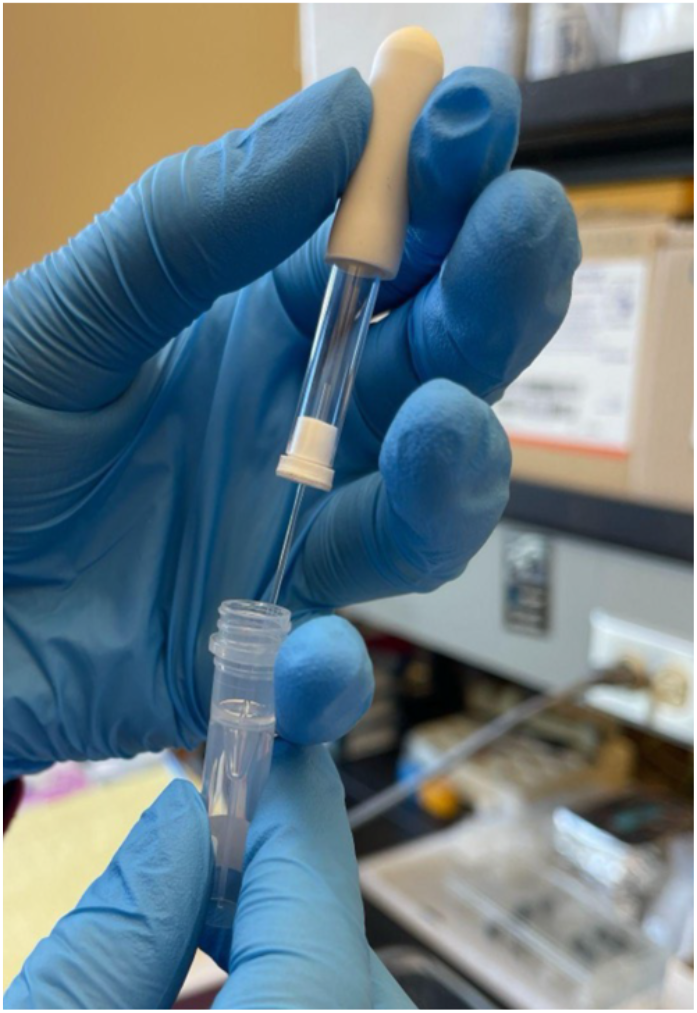
Transference of Collected CSF. Using a rubber bulb capillary dispenser into a storage vial in a laboratory setting. This process ensures the integrity and sterility of the sample for subsequent analyses.

### 2.5 Brain Extraction

1. Place the mouse in a ventral position.
2. Spray the cranium with 70% EtOH to wet the hair down.
3. Using a sharp pair of large scissors cut transversely immediately caudal to the occiput. Discard the body appropriately. *If removal of the head from the body is desired, follow this step. If not, go to step 4.*
4. Incise the cutaneous layers using the small scissors in a midline sagittal cut extending from caudal occiput to the rostrum, see **Figure 7A**. Displace the skin tissue laterally to visualize the skull.
5. Place the tips of the small scissors in the orbitals and cut the skull transversely as seen in **Figure 7B**.
6. Fixing the tips of the small scissors under the occiput of the skull, with slow rostral progression, cut on midline with small precise movements to the previous orbit incision, see **Figure 7C**.
7. Placing the forceps under the skull at the incision on the left side, grip and pull the skull laterally, perpendicular to the midline, peeling the skull from the brain, **see Figure 7D**.
8. If the skull breaks into pieces, remove each piece individually until complete hemisphere exposure is achieved.
9. Repeat steps 7 and 8 on the right hemisphere.
10. Once full brain exposure is accomplished, place the forceps beneath the brainstem and excise the brain by gently pulling superiorly and rostrally. *Inverting the mouse so that the dorsal surface of the cranium is closest to the workbench can be helpful, as it allows gravity to assist in brain extraction*.

**Figure 7.**
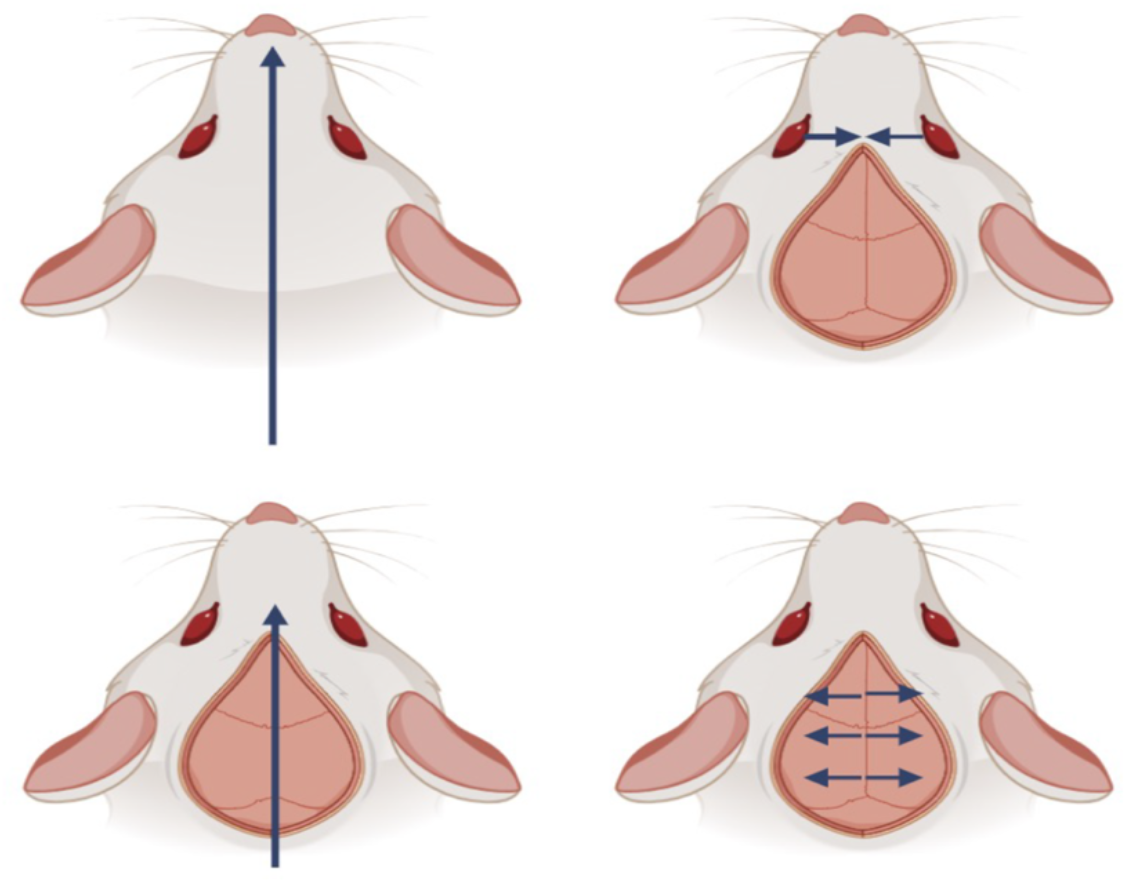
Brain Extraction. **[A]** Midline sagittal skin incision from occiput to rostrum. **[B]** Transverse incision between the orbitals. **[C]** Skull cut from occiput to orbit in a rostral direction. **[D]** Lateral separation of skull from brain. Created in BioRender. Seerley, A. (2025) https://BioRender.com/k5txf1t

### 2.6 Brain Microdissection

The following procedure was performed under a microdissection microscope. Full dissection shown in **Figure 8.** See **Figure 9** for a size reference of the mouse brain hemisphere after separation.

1. *Hemispheric Separation*—Place the brain on a clean surface with the dorsal side facing up (ventral side facing down) **(Figure 10A).** Using the scalpel, make a sagittal cut along the longitudinal fissure to separate the hemispheres.
2. *Thalamus*—On the medial side of the brain while holding the hemisphere steady with forceps, visualize and scoop out the thalamus with the scalpel using blunt dissection paying attention to region boundaries, see **Figure 10B**.
3. *Hippocampus*— On the medial side of the brain, peel the cortex laterally, perpendicular to the midline using blunt dissection with the scalpel and while holding the hippocampus in place with the forceps to ensure separation of the two regions. Perform blunt dissection to extract the hippocampus medially toward the midline using the scalpel, see **Figure 10C and D**.
4. *Striatum*—Expose the striatum by removing the cortex and hippocampus as described above. Incise around the perimeter of the striatum with fine tip thumb forceps. Using the scalpel scoop out the striatum with reference to natural tissue separation, see **Figure 10E and F**. Striations help to differentiate the striatum from the cortex.
5. *Cortex*—Excise the base of the already dissected cortex using the scalpel blade, see **Figure 10G**.
6. *Cerebellum*—Remove the base of the cerebellum from the rest of the brain using a scalpel with attention to natural separation, see **Figure 10H**.
7. *Brainstem*—Excise the brain stem from the midbrain, see **Figure 10I**.
8. *Olfactory bulb*—Excise the olfactory bulb from the forebrain, see **Figure 10J**.

**Figure 8.**
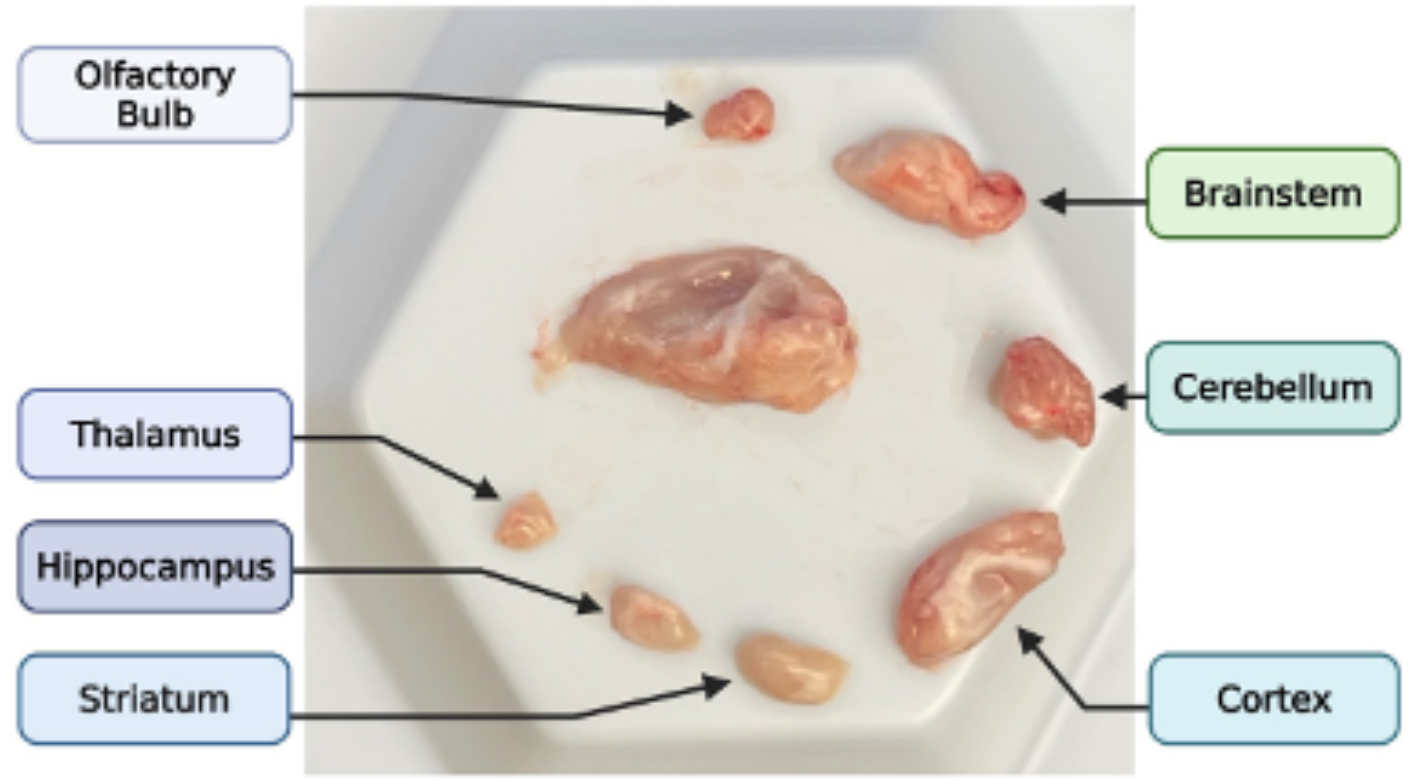
Completed Microdissection of the Mouse Brain. Labeled brain regions following careful dissection including the olfactory bulb, thalamus, hippocampus, striatum, brainstem, cerebellum, and cortex. Created in BioRender. Seerley, A. (2025) https://BioRender.com/v5y4pwe

**Figure 9.**
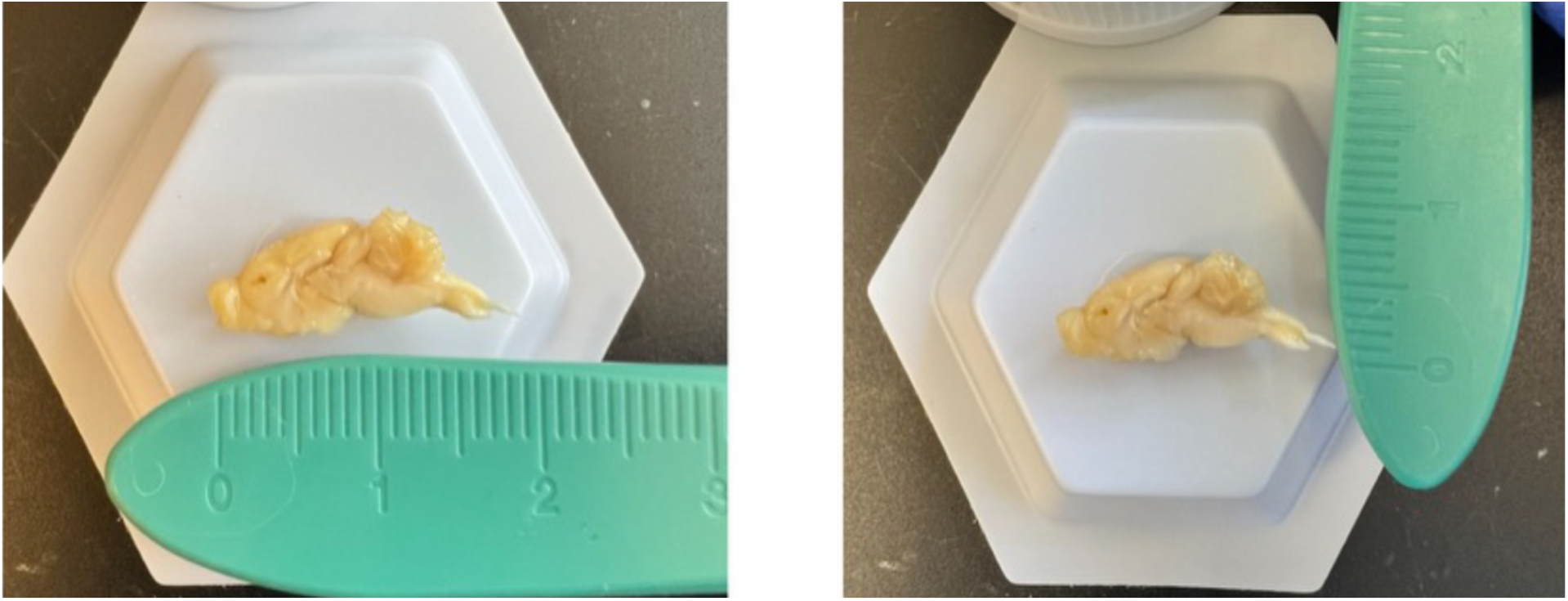
Size Reference of the Mouse Brain. Lateral-medial view of a fixed right hemisphere mouse brain. **[A]** Measuring approximately ∼1.9cm in length and **[B)**]∼0.6cm in height.

**Figure 10.**
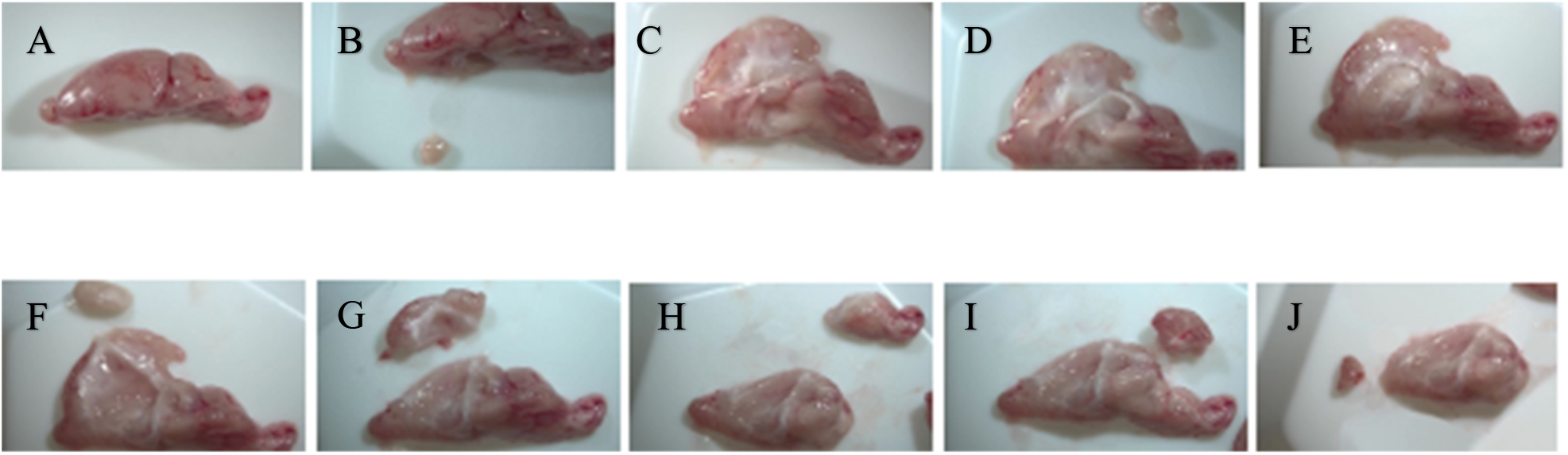
Brain Microdissection. Panel **[A]** shows the right hemisphere. **[B]** Thalamus, **[C&D]** Hippocampus, **[E&F]** Striatum, **[G]** Cortex, **[H]** Cerebellum, **[I]** Brainstem, **[J]** Olfactory bulb. (Zeiss Stemi 508 & Axiocam 208 Color Photo)

### 2.7 Sciatic Nerve Dissection

1. Place the mouse on the desired side up and pin the feet to the display tray.
2. Remove the skin over the quadriceps exposing the lateral surface.
3. Visualize the sciatic nerve and do not mistake the femur or muscle fascia for the sciatic nerve, see **Figure 11**.
4. Open surrounding muscle layers, following natural tissue separation, see **Figure 11**.
5. Once exposed, leaving nerve in place, fill the space with formalin for 30 minutes if fixation is desired. Make sure to not stretch or disturb the nerve as the sciatic nerve tissue is extremely delicate and easily damaged.
6. After the nerve has been fixed, cut the nerve at the head of femur down to the patella region.
7. Remove the nerve and preserve as desired.

**Figure 11.**
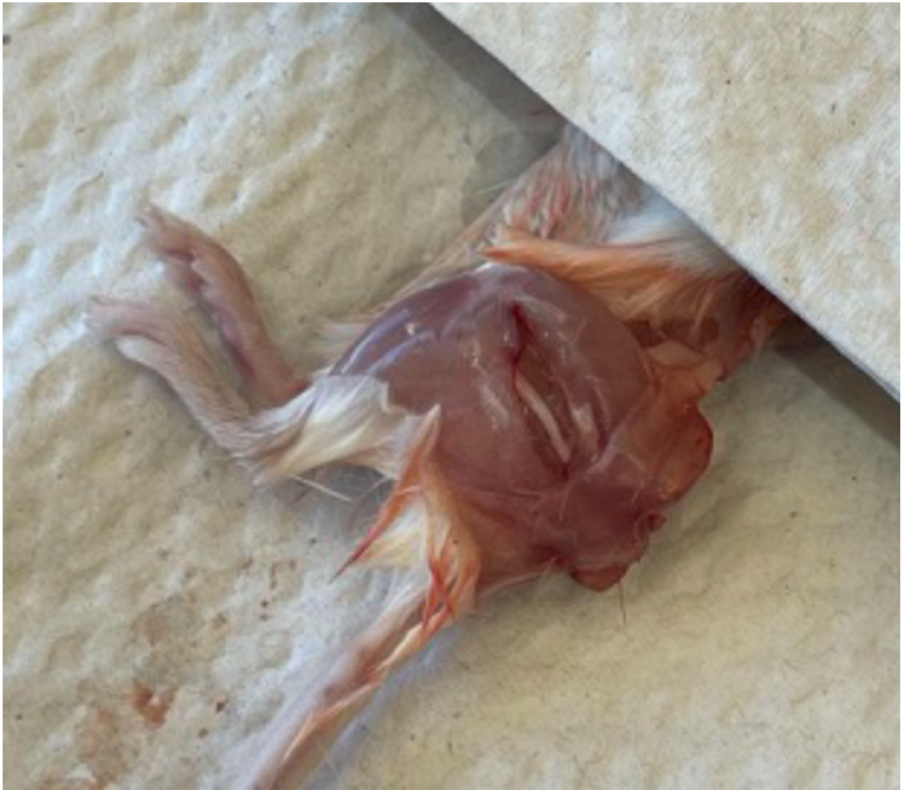
Proper positioning of an Anesthetized Mouse for Sciatic Nerve Extraction. The mouse is placed on its side with its hind limbs extended and secured to expose the sciatic nerve pathway, ensuring accurate and efficient nerve dissection.

### 2.8 Spinal Cord Extrusion

1. Place mouse on ventrum and coat with EtOH.
2. Decapitate the mouse at the occiput.
3. Remove the skin covering the spine.
4. Starting at the cranial side of the body using the large scissors, slice through the tissues and ribs lateral to the spinal vertebrae progressing caudally to the pelvis.
5. Next, perform a transverse scissor slice immediately cranial to the pelvis. This should extract the spine from the cervical vertebrae to the lumbosacral junction.
6. Remove additional organs with forceps and scissors on the ventral side of the spine.
7. Starting at the caudal end, ensure that the spinal cord is clean by removing excess tissue if necessary.
8. Hold the spinal cord in a fashion that eliminates the curvature.
9. Using a 10ml syringe filled with phosphate buffered saline (PBS), secure a tight seal with the syringe (may need to add a pipette tip with the tip cut to fit the size of the spinal cord) to the spinal cord cavity, allowing hydrostatic pressure to extrude the spinal cord into a petri dish filled with phosphate buffered saline (PBS), see **Figure 12A**. The extrusion of spinal cord may be easier if specific spinal regions are dissected. This can be accomplished with transverse scissor slices at the thoracolumbar junction to isolate the thoracic spinal cord from the lumbar spinal cord, see **Figure 12B and C**. If the spinal cord is stuck to the end of the vertebrae, carefully use the forceps to manipulate the spinal cord out of the cavity.
10. Store as desired.

**Figure 12.**
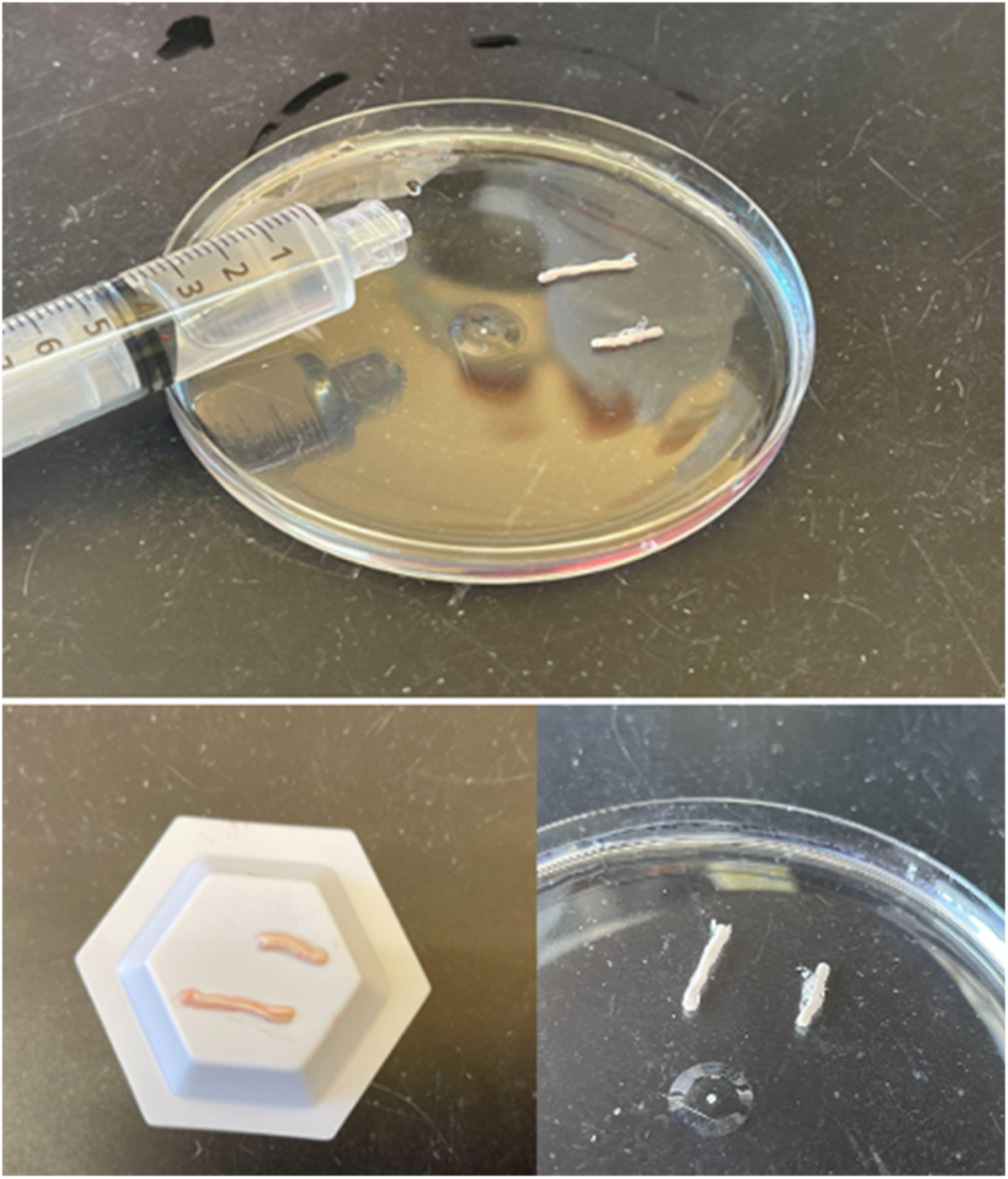
Spinal Cord Post-Extraction. **[A]** Thoracic and lumbar spinal cord regions in petri dish after extrusion with a 10ml syringe **[B, C]** View of the thoracic and lumbar spinal cord regions after extrusion; ready for further processing and analysis.

## 3. Results

### 3.1 Applications of CSF in Neurodegenerative Disease Studies

CSF is the fluid surrounding the central nervous system (CNS), providing needed nutrients and removing waste products. Analysis of the CSF gives real-time insight into the biochemical state of the CNS, such as biomarker detection, disease progression tracking, and pharmaceutical testing (Kaur *et al.*, 2023). In mice and most rodents, the cisterna magna is the typical site for CSF collection due to its easier accessibility and large volume (Liu and Duff, 2008). This fluid collection is integral in translational research, so proper handling and curation are required to have reputable results (Liu and Duff, 2008; Lim *et al.*, 2018; Shimizu *et al.*, 2022; Kaur *et al.*, 2023).

### 3.2 Applications of Brain Extraction in Neurodegenerative Disease Studies

Dissection of the whole brain is an essential technique to acquire tissue for many methods in neurodegenerative disease studies. Among these methods, histology is a commonly used to analyze basic morphology and the quantity of specific cell types or proteins of interest, requiring precise extraction to maintain tissue integrity. Histologic examples of brain hemispheres are shown displaying spongiform change surrounding the dentate gyrus of the hippocampus in transgenic mice infected with a prion disease **(Figure 13A)** when compared to a control **(Figure 13B).**

**Figure 13.**
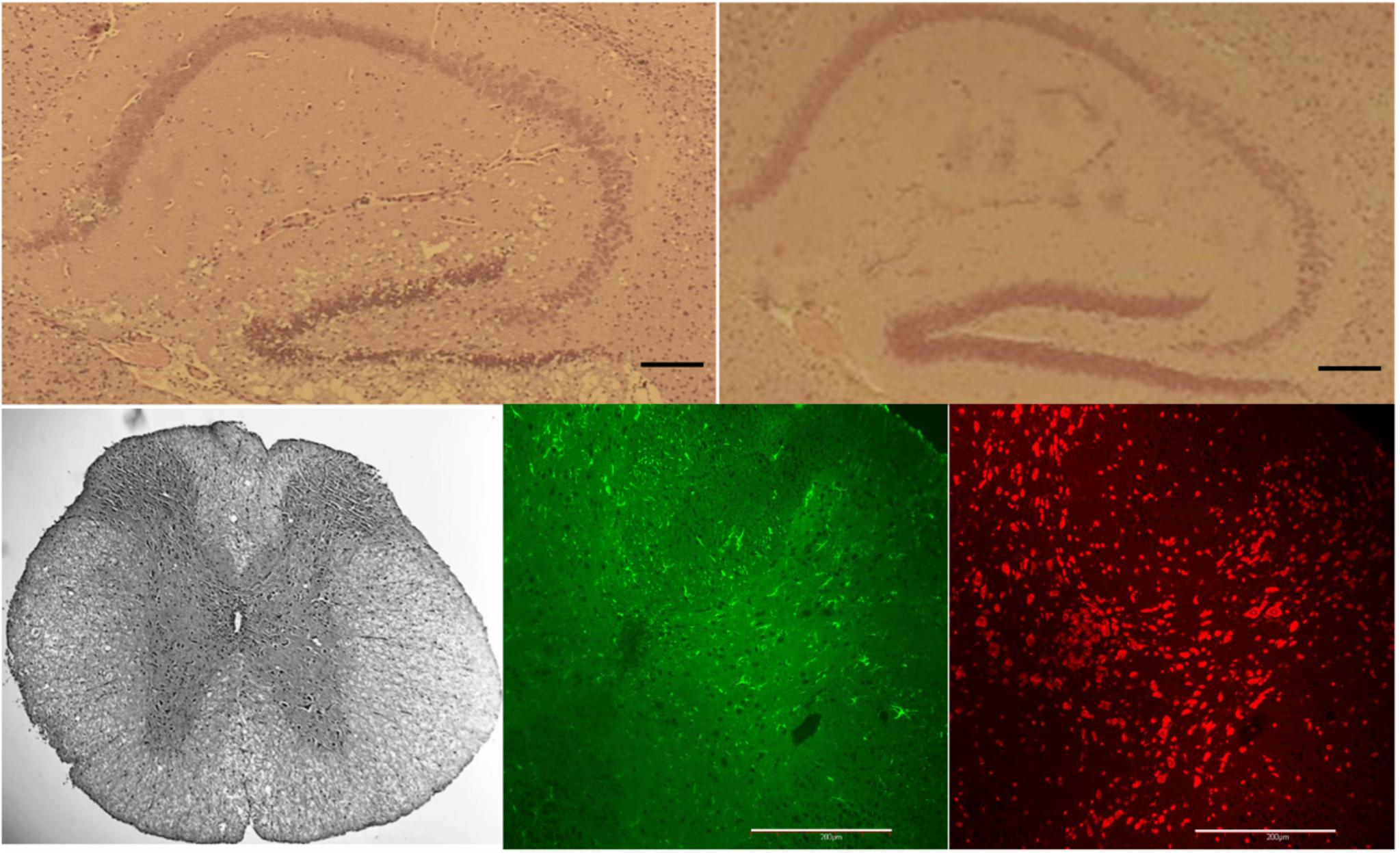
Examples of histologic preparations after brain extraction and spinal cord extrusion demonstrating the importance of maintaining tissue integrity for histologic studies. **[A]** Chronic wasting disease (CWD) transgenic mouse model brain hemisphere sections processed and stained with hematoxylin and eosin (H&E) exhibiting spongiform tissue around the dentate gyrus of the hippocampus as opposed to non-diseased control brain in panel **[B].** Scale bar 1mm. **[C]** Spinal cord section processed and stained with H&E to display spinal cord morphology, shown in black and white. **[D]** Presence of astrocytes (green) in spinal cord sections after immunofluorescence (IF) staining in CWD mouse models, scale bar 200um. **[E]** Spinal cord section of IF neuronal labeling (red), scale bar 200um.

### 3.3 Applications of Microdissected Brain Regions in Neurodegenerative Disease Studies

Often, neurodegenerative disease studies require dissection of specific brain regions due to specific region neuropathology. For example, prion diseases cause morphological changes in the hippocampus and cortex regions of the brain (Bonda *et al.*, 2016; Orge *et al.*, 2021), which require precise isolation for use in research. In relation to Huntington’s disease, studies are typically focused on the striatum as it is the main region affected (Ament *et al.*, 2017; Bragg *et al.*, 2024). Similarly, biomarker research in neurological studies is most accurate and reproducible when conducted on distinct sections of the brain because it eliminates data dilution from regions that may have less pathology. These methods are also important in pharmacology development studies as brain regions are often required to better understand pharmaceutical drug distribution, safety, and mechanisms of action (Zalachoras *et al.*, 2013).

### 3.4 Applications of Sciatic Nerve in Neurodegenerative Disease Studies

The largest and longest nerve in the rodent and human body is the sciatic nerve. It extends from the pelvic region and tracks along the hind limbs. Due to its size and simple accessibility, this peripheral nerve is integral to nerve studies. As it is part of the peripheral nervous system (PNS), it can give information about peripheral neuropathies, neuromuscular junction processes, therapeutics, and Wallerian degeneration (Toma, McPhail and Ramer, 2007; Bala *et al.*, 2014; Sell, Shi and Bhat, 2024).

### 3.5 Applications of Spinal Cord in Neurodegenerative Disease Studies

The dynamic conduit for connection between the brain and body of sensory and motor signaling is the latter half of the CNS, the spinal cord. Spinal cord research gives insight into motor neuron diseases, degeneration and regeneration processes, inflammation, and traumatic injuries. Extrusion of the spinal cord is commonly performed via hydrostatic pressure, which allows gentle expulsion without tissue disturbance or contamination (Richner *et al.*, 2017). An example of this is shown **(Figure 13C)** with hematoxylin and eosin staining of spinal cord sections using this extrusion protocol. Immunofluorescent staining displays neurons (in red) and astrogliosis (in green) in cervidized transgenic mice infected with chronic wasting disease (CWD), a fatal prion disease found in cervids **(Figure 13D and E).**

### 3.6 Dissection Troubleshooting

Reproducibility is crucial for scientific studies; however, there are many errors that may occur when performing dissections such as the ones listed above. It’s important to recognize potential mistakes and avoid them when possible. Common errors and solutions are addressed in **Table 1**.

**Table 1.**
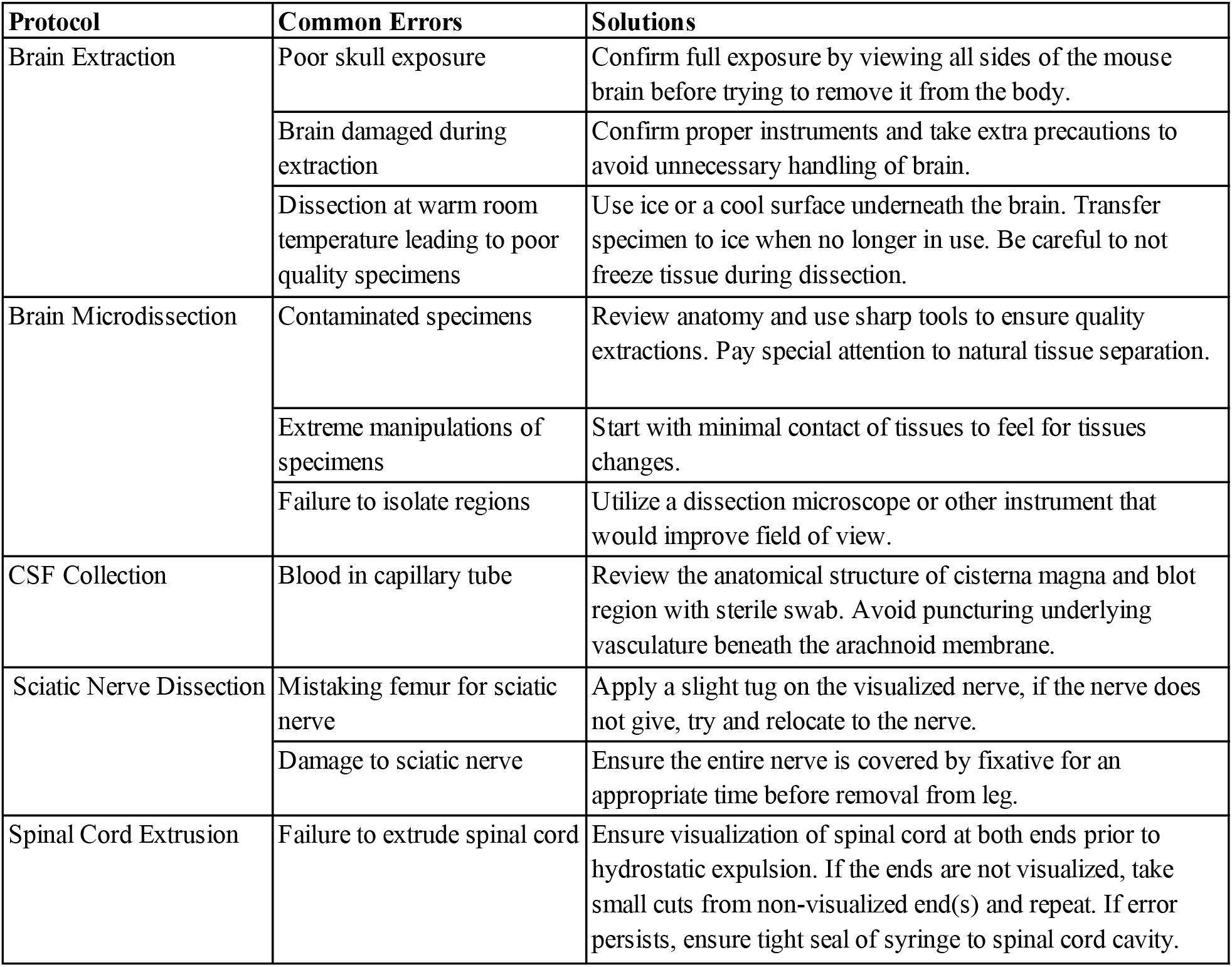
Common errors and troubleshooting solutions arranged by protocol.

## 4. Discussion

High-quality, reproducible dissection protocol techniques are the backbone of experimental neuroscience. Whether the focus is on the CNS or PNS, consistency ensures that biological variability drives the observed outcomes, not technical error (Voelkl *et al.*, 2018). These protocols maintain a high level of tissue integrity that allows for cross-laboratory collaborative comparison, which is a necessity in translational research (Richner *et al.*, 2017; Meyerhoff *et al.*, 2021). Optimization also reduces animal usage, maximizes data per specimen, and minimizes overall suffering. As neuroscience research continues to intersect with technology, clinical medicine, and ethics, standardized protocols are called to transcend previous frameworks/compositions and establish new benchmarks for precision, reproducibility, and ethical rigor (Krakauer *et al.*, 2017).

Meticulous dissection of the brain and nervous system tissue is fundamental in reproducible, top-caliber neurodegenerative research. Accurate replication of dissection protocols ensures that key anatomical regions can be isolated without contamination or damage to the tissue’s integrity. Detailed protocols like the ones described in this paper successfully provide researchers with ideal samples that can be used in a vast number of applications. For example, prion diseases are found to be concentrated in the hippocampus and cortex (Bonda *et al.*, 2016; Orge *et al.*, 2021). Isolation of these brain regions in prion disease studies prevents biomarker dilution and enables targeted histopathological analysis. In relation to Huntington’s disease, proteomic transcriptomic, and pharmaceutical studies are conducted on the striatum as it is the main region affected by neurodegeneration (Ament *et al.*, 2017; Bragg *et al.*, 2024). Brain microdissections are often required in pharmaceutical research studies such as investigations of antisense oligonucleotide (ASO) therapies to better understand drug distribution, safety profiles, and pharmacodynamics.

These protocols lay the groundwork for translational advances and bridge laboratory practices to clinical skills. Precise anatomical knowledge directly translates into surgical practices where understanding the location and relationships of structures guides every move. Dissections like these may even be useful for training future physicians by developing fine motor technical skills with refined surgical tools, which are required in microsurgery settings (Kshettry *et al.*, 2014). Performing brain microdissections under the microscope also builds a better understanding of how to work with varied depth-of-field, similar to robotic surgery skills that occur under magnification, enhancing operative medicine. Lastly, adherence to biosafety protocols (BSL-2) during dissections mirrors the sterile techniques of surgical suites, not only reinforcing good habits but once again preventing contamination. These practices protect both the research and tissue integrity to model the necessary standards for plausible and reputable research findings.

## 5. Conclusion

In closing, implementation of standardized protocols across the brain, spinal cord, CSF, and peripheral nerves such as these provided, allows for maximized tissue specimen value, reduced animal usage, and more robust experimental power. Furthermore, it encourages cross disciplinary benefits in pharmaceutical, forensic, bioengineering, and regenerative medicine, allowing for a broader impact.

## Data Availability Statement

All methods and data described in this manuscript will be freely accessible and available to any researcher wishing to use them.

## Ethics Statement

All experiments were conducted under approved Institutional Animal Care and Use Committee protocol (McLaughlin Research Institute IACUC 2023-AG-100).

## Author Contributions

LC: Conceptualization, Visualization, Data curation, Data presentation, Writing-original draft, Writing-review & editing. AS: Product administration, Resources, Data curation, Writing-original draft, Writing-review & editing. SE: Supervision, Resources, Data curation, Writing-review & editing. AGP: Conceptualization, Visualization, Methodology, Funding acquisition, Supervision, Validation, Writing-original draft, Writing-review & editing.

## Acknowledgments

Colony management for the mouse models included in this study was provided by the McLaughlin Research Institute -Gene Editing and Mouse Models Assessment (GEMMA) Core Facility within the Center for Integrated Biomedical and Rural Health Research, RRID:SCR_027045, 1P20GM152335. The authors thank Rose Pitstick and Kaela Davies for exceptional care of the mice. We would also like to thank student intern research assistant Bridget Gray and federal work study Touro COM-MT students Dalia Shaaban and Clairissa Kaylor for thoughtful discussions regarding planning and presentation of dissection methods in the mouse brain microdissections.

## References

Aboghazleh, R. et al.. (2024) ‘Rodent brain extraction and dissection: A comprehensive approach’, MethodsX, 12, p. 102516. Available at: 10.1016/j.mex.2023.102516.

Ament, S.A. et al.. (2017) ‘High resolution time-course mapping of early transcriptomic, molecular and cellular phenotypes in Huntington’s disease CAG knock-in mice across multiple genetic backgrounds’, Human Molecular Genetics, 26(5), pp. 913–922. Available at: 10.1093/hmg/ddx006.

Bala, U. et al.. (2014) ‘Harvesting the maximum length of sciatic nerve from adult mice: a stepby-step approach’, BMC research notes, 7, p. 714. Available at: 10.1186/17560500-7-714.

Bonda, D.J. et al.. (2016) ‘Human prion diseases: surgical lessons learned from iatrogenic prion transmission’, Neurosurgical Focus, 41(1), p. E10. Available at: 10.3171/2016.5.FOCUS15126.

Bondulich, M.K. et al.. (2024) ‘Translatable plasma and CSF biomarkers for use in mouse models of Huntington’s disease’, Brain Communications, 6(1), p. fcae030. Available at: 10.1093/braincomms/fcae030.

Bragg, R.M. et al.. (2024) ‘Global huntingtin knockout in adult mice leads to fatal neurodegeneration that spares the pancreas’, Life Science Alliance, 7(9), p. e202402571. Available at: 10.26508/lsa.202402571.

Chinwalla, A.T. et al.. (2002) ‘Initial sequencing and comparative analysis of the mouse genome’, Nature, 420(6915), pp. 520–562. Available at: 10.1038/nature01262.

Cook, M., Hensley-McBain, T. and Grindeland, A. (2023) ‘Mouse models of chronic wasting disease: A review’, Frontiers in Virology, 3. Available at: 10.3389/fviro.2023.1055487.

Kaur, A. et al.. (2023) ‘A protocol for collection and infusion of cerebrospinal fluid in mice’, STAR protocols, 4(1), p. 102015. Available at: 10.1016/j.xpro.2022.102015.

Krakauer, J.W. et al.. (2017) ‘Neuroscience Needs Behavior: Correcting a Reductionist Bias’, Neuron, 93(3), pp. 480–490. Available at: 10.1016/j.neuron.2016.12.041.

Kshettry, V.R. et al.. (2014) ‘The role of laboratory dissection training in neurosurgical residency: results of a national survey’, World Neurosurgery, 82(5), pp. 554–559. Available at: 10.1016/j.wneu.2014.05.028.

Kuttner-Hirshler, Y. et al.. (2017) ‘Brain Biomarkers in Familial Alzheimer’s Disease Mouse Models’, Journal of Alzheimer’s disease : JAD, 60(3), pp. 949–958. Available at: 10.3233/JAD-170269.

Lim, N.K.-H. et al.. (2018) ‘An Improved Method for Collection of Cerebrospinal Fluid from Anesthetized Mice’, Journal of Visualized Experiments: JoVE, (133), p. 56774. Available at: 10.3791/56774.

Liu, L. and Duff, K. (2008) ‘A Technique for Serial Collection of Cerebrospinal Fluid from the Cisterna Magna in Mouse’, JoVE (Journal of Visualized Experiments), (21), p. e960. Available at: 10.3791/960.

Meyerhoff, J. et al.. (2021) ‘Microdissection of Mouse Brain into Functionally and Anatomically Different Regions’, Journal of Visualized Experiments: JoVE [Preprint], (168). Available at: 10.3791/61941.

Minikel, E.V. et al.. (2020) ‘Prion protein lowering is a disease-modifying therapy across prion disease stages, strains and endpoints’, Nucleic Acids Research, 48(19), pp. 10615–10631. Available at: 10.1093/nar/gkaa616.

Orge, L. et al.. (2021) ‘Neuropathology of Animal Prion Diseases’, Biomolecules, 11(3), p. 466. Available at: 10.3390/biom11030466.

Ratz-Mitchem, M.L. et al.. (2023) ‘Generation and characterization of a knock-in mouse model for spastic tetraplegia, thin corpus callosum, and progressive microcephaly (SPATCCM)’, Mammalian Genome: Official Journal of the International Mammalian Genome Society, 34(4), pp. 572–585. Available at: 10.1007/s00335-023-10013-4.

Richner, M. et al.. (2017) ‘Hydraulic Extrusion of the Spinal Cord and Isolation of Dorsal Root Ganglia in Rodents’, Journal of Visualized Experiments : JoVE, (119), p. 55226. Available at: 10.3791/55226.

Sell, L.B., Shi, Q. and Bhat, M.A. (2024) ‘Protocol for isolating and processing mouse sciatic nerve fibers for confocal immunohistochemistry’, STAR Protocols, 5(1), p. 102852. Available at: 10.1016/j.xpro.2024.102852.

Shimizu, K. et al.. (2022) ‘Anatomy-oriented stereotactic approach to cerebrospinal fluid collection in mice’, Brain Research, 1774, p. 147706. Available at: 10.1016/j.brainres.2021.147706.

Sultan, F.A. (2013) ‘Dissection of Different Areas from Mouse Hippocampus’, Bio-Protocol, 3(21), p. e955. Available at: 10.21769/bioprotoc.955.

Tello, J.A. et al.. (2022) ‘Animal Models of Neurodegenerative Disease: Recent Advances in Fly Highlight Innovative Approaches to Drug Discovery’, Frontiers in Molecular Neuroscience, 15. Available at: 10.3389/fnmol.2022.883358.

Toma, J.S., McPhail, L.T. and Ramer, M.S. (2007) ‘Differential RIP antigen (CNPase) expression in peripheral ensheathing glia’, Brain Research, 1137(1), pp. 1–10. Available at: 10.1016/j.brainres.2006.12.053.

Vallabh, S.M. et al.. (2023) ‘Therapeutic Trial of anle138b in Mouse Models of Genetic Prion Disease’, Journal of Virology, 97(2), p. e0167222. Available at: 10.1128/jvi.01672-22.

Voelkl, B. et al.. (2018) ‘Reproducibility of preclinical animal research improves with heterogeneity of study samples’, PLoS biology, 16(2), p. e2003693. Available at: 10.1371/journal.pbio.2003693.

Young, A.B. (2009) ‘Four Decades of Neurodegenerative Disease Research: How Far We Have Come!’, The Journal of Neuroscience, 29(41), pp. 12722–12728. Available at: 10.1523/JNEUROSCI.3767-09.2009.

Zalachoras, I. et al.. (2013) ‘Antisense-mediated isoform switching of steroid receptor coactivator-1 in the central nucleus of the amygdala of the mouse brain’, BMC Neuroscience, 14(1), p. 5. Available at: 10.1186/1471-2202-14-5.

